# A common set of structurally distinct endogenous retroviruses predict survival in multiple human tumor types

**DOI:** 10.1101/2024.02.07.579350

**Authors:** Alexandra L. Massa, Himanshu Chintalapudi, Evelyn C. Coppenbarger, Jake M. Peterson, Walter N. Moss, H. Robert Frost, Steven D. Leach

**Affiliations:** Department of Molecular and Systems Biology and the Dartmouth Cancer Center, Geisel School of Medicine at Dartmouth, Lebanon, New Hampshire; Department of Biomedical Data Science, Geisel School of Medicine at Dartmouth, Lebanon, New Hampshire; Roy J. Carver Department of Biophysics, Biochemistry and Molecular Biology, Iowa State University, Ames, Iowa

## Abstract

Accumulating evidence suggests that viral mimicry induction in cancer cells through the expression of human endogenous retroviral elements (ERVs) enhances tumor immunogenicity and improves response to immunotherapy. However, the characterization of ERV expression across multiple tumor types, their clinical relevance, and their role in tumor immunity remain limited. Here, we assessed the prognostic and immune-related consequences of ERV expression across 8 different tumor types from The Cancer Genome Atlas (TCGA). We applied a recently developed tool, Telescope, to quantify the expression of 14,968 ERVs in bulk tumor RNA-sequencing data. Approximately 15-22% of Telescope’s ERVs (n_avg_ = 2,736) were expressed in each tumor cohort. We identified both tissue-specific and commonly expressed ERVs across cohorts. Over 50% of tissue-specific ERVs were ubiquitously expressed (n = 1,709) across all 8 tumor types. Using patient clinical data, we identified a subset of 94 ERVs whose expression levels were significantly associated with overall survival, of which 42 were favorably prognostic. Prognostic ERVs displayed unique structural characteristics, including more frequent inverted repeats and more stable secondary structures. Patients were classified based on ERV expression, allowing identification of patient cohorts with greater than 6-fold differences in overall survival. We also found that expression of positively prognostic ERVs was associated with enrichment for selected anti-tumor immune response gene expression signatures. This analysis extends previous results identifying patterns of ERV expression in bulk tumor RNA-seq datasets, providing further insight into the role of ERVs in tumor clinicopathology and immunogenicity.

## Introduction

Human endogenous retroviruses (ERVs) are remnant sequences of past exogenous retroviruses that infected our ancestral germline millions of years ago. Strikingly, ERVs comprise ∼8% of our genome – nearly four times that of protein-coding sequences – and exist as a permanent, ubiquitous part of our DNA [1]. Their nascent gene structures resemble exogenous retroviruses, containing essential viral protein-coding genes flanked by 5′ and 3′ regulatory regions known as long terminal repeats (LTRs). However, due to accumulating mutations and deletions throughout evolution, most ERVs exist as fragmented, non-infectious solitary LTRs or partial open reading frames (ORFs). In healthy tissue, most ERV sequences are transcriptionally silenced by strict epigenetic mechanisms; however, in diseased states such as cancer, ERVs may become reactivated.

Evidence suggests ERV expression may induce viral mimicry signals in cancer cells, associated with enhanced immunogenicity and improved patient response to immunotherapy [2–15]. Studies have shown a positive association between ERV transcript levels and innate immune activation, resembling the response against viral double-stranded RNA (dsRNA) often observed in virus-infected cells [4,5,9,13]. Derepressed LTR sequences of ERVs, which retain transcriptional regulatory function, are functionalized as enhancers for host interferon (IFN) gene networks and can also serve as alternative promoters generating truncated or chimeric proteins with immunogenic potential [13,16]. Indeed, increased ERV transcript levels correlate with immune checkpoint response in urothelial and renal cell cancers, positioning ERVs as an alternative source of tumor-associated antigens [10,12].

Despite these findings, the characterization of ERV expression across multiple tumor types, the determination of their clinical relevance, and their more general role in tumor immunity remain incomplete. This fragmented knowledge is due to 1) a historically small reference set of ERV transcripts to query *in silico* and 2) the repetitive nature of ERV sequences, which complicates genomic mapping. Over the last several years, there have been significant advances in the annotation of ERVs and the ability to map these repetitive sequences in RNA-sequencing (RNA-seq) data [17]. Yet nearly all these studies evaluate bulk, global expression of ERVs rather than pinpointing the exact location of a specific sequence.

Here, we apply a recently developed bioinformatic tool, *Telescope* [18], to quantify the expression of 14,968 ERVs at specific genomic locations across 8 types of tumors using publicly available data from The Cancer Genome Atlas (TCGA). We performed survival analysis based on overall patient survival and correlative analysis against the expression of immunologically relevant gene sets. This allowed us to examine the prognostic and immune-related consequences of ERV expression in various human malignancies.

## Materials and methods

### TCGA datasets

The utilization of TCGA data (accession phs000178.v11.p8) was authorized by the National Cancer Institute (NCI) data project access committee through the Database of Genotypes and Phenotypes (dbGaP; https://dbgap.ncbi.nlm.nih.gov) on June 15, 2020. Paired-end RNA-seq data provided by TCGA were downloaded and analyzed between June 15, 2020-June 15, 2023. BAM-formatted read alignment files (for GRCh38/hg38) of the RNA-seq data were downloaded from the Genomic Data Commons (GDC) data portal site (https://portal.gdc.cancer.gov/) using the command-line tool gdc-client v1.5.0 (https://gdc.cancer.gov/access-data/gdc-data-transfer-tool/) for a total of 8 cohorts (TCGA-PAAD, TCGA-LIHC, TCGA-KIRC, TCGA-COAD, TCGA-OV, TCGA-BLCA, TCGA-LUAD, TCGA-BRCA). Only treatment-naïve samples of the primary tumor type (non-metastatic) were considered; all recurrent/metastatic tumor samples and solid tissue normal samples were excluded. If patient samples had multiple tumor samples, sample IDs with only one vial designation were included; if patient samples had multiple plate identifiers, the sample with the highest lexicographical sort value of the plate was considered. In total, samples from 135 pancreatic cancer (PAAD) cases, 342 liver cancer (LIHC) cases, 463 kidney cancer (KIRC) cases, 321 colorectal cancer (COAD) cases, 365 ovarian cancer (OV) cases, 390 bladder cancer (BLCA) cases, 481 lung cancer (LUAD) cases, and 445 breast cancer (BRCA) cases were used for downstream analysis (S Table 1). Corresponding publicly available clinical data, including tumor stage, grade, and survival status, were also downloaded. No identifying information for individual patients was available during or after data collection.

### ERV quantification and characterization

BAM-formatted alignment files were name-sorted using samtools version 1.9 and converted to fastq format using samtools command bam2fq with parameters to avoid secondary alignments (-F 0×100) or bedtools v2.26.0 using bamtofastq command. Fastq files were aligned to the human genome (hg38) using bowtie2 (version 2.3.4.3) using the following parameters: suppress SAM records for unaligned reads (--no-unal), local alignment search (--very-sensitive-local), allow up to 100 alignments per read (-k 100), minimum alignment score (--score-min L,0,1.6). Bowtie2 aligned BAM files were subjected to the ERV quantification pipeline Telescope version 1.0 using the assigned module and the ERV annotation for the hg38 genome provided (https://github.com/mlbendall/telescope_annotation_db) with --theta_prior of 200000 and --max_iter 200 and utilizing custom scripts [18]. The output generated by Telescope is a table of ERVs (labeled by chromosomal location) and their relative expression, quantified by read counts. Final counts from the output were loaded into DESeq2 using the DESeqDataSetFromMatrix function [19]. Normalization size factors were calculated by dividing the library sizes by their geometric mean. Variance stabilizing transformation was applied using DESeq2’s vst function, and the normalized counts were used for subsequent analysis. ERVs with an expression cut-off of the sum of 10 raw counts in at least 50% of samples were considered as expressed ERVs in each cohort, and the rest were excluded from any analysis. ERV family level expression was calculated by summing the final counts from Telescope across the family level from the family level annotation provided here: https://github.com/mlbendall/telescope_annotation_db/tree/master/refs. Genomic meta-features of locus-specific ERVs were obtained from https://github.com/Liniguez/Telescope_MetaAnnotations. Protein-coding potential of ERV elements was calculated based on intrinsic sequence composition using the Coding-Noncoding-Identifying Tool (CNIT)[20]. Inverted repeats were identified using the Inverted Repeats Finder (IRF) Software [21].

### Transcriptome quantification

Gene expression was quantified using STAR aligner (v.2.7.3a) run on fastq files using the STAR genome index generated from the human genome hg38 (GRCh38.p12) fasta file and GENCODE (v.31) gtf file. Read counting was done in unstranded mode using featureCounts (v.1.6.4). To normalize read counts, either variance stabilizing transformation (vst) was applied using DESeq2’s vst function or transcripts per million (TPM) counts were generated using gene lengths calculated from the sum of the union of exonic sizes from the gtf file. Normalized read counts were used in all subsequent analyses.

### Survival analysis

Survival analysis was performed in R using the package survival (v.3.2-7) (https://cran.r-project.org/web/packages/survival/index.html). Clinical metadata of survival time, vital status, age, tumor grade, and tumor stage were downloaded from the GDC data portal. Survival time in days or months was obtained from fields *death_days_to* and *last_contact_days_to*; if survival time was not available for *death_days_to*, *last_contact_days_to* was considered, and vice-versa. Patients with unavailable survival time or values <0 were removed from the analysis. Patients with tumor stage having values like “Discrepancy” or “Not Available” in fields like *ajcc_pathologic_tumor_stage* were removed from the analysis, and any sub-stages in *ajcc_pathologic_tumor_stage* were consolidated to the main stages I, II, III, IV. Censoring values for vital status were encoded as 0 for alive and 1 for dead. Univariate Cox regression was performed with the Coxph function using the R package RegParallel for parallelization (https://github.com/kevinblighe/RegParallel). A Cox proportional hazards regression model was fit using patients’ overall survival and each ERV’s variance-stabilized expression value as a continuous variable while adjusting for patient age, tumor stage, and tumor grade. Only age and tumor stage were considered for cohorts where tumor grade was not present or obtainable.

To estimate the impact of ERV expression on survival from all the cohorts, 1709 intersecting normalized ERV counts from all 8 cohorts were pooled, and glmnet with LASSO-penalized regression was applied using the R package glmnet (https://cran.r-project.org/package=glmnet). Nested cross-validation was used to train and evaluate a Lasso-penalized Cox proportional hazards model using the 1,709 ERVs. Cross-validation (CV) was employed both for the training versus test split (5-fold CV) and within each training fold (10-fold CV) for selection of the Lasso penalty parameter. The mean of the means of λ (lambda.min) was chosen with the non-zero coefficient estimates and their predictor ERVs. With the 449 selected ERVs and estimated parameters, we ran our prediction on the test data by applying the Cox proportional hazards regression model identified in the training data. 94 ERVs were significant at FDR-adjusted p<0.05, with 42 ERVs with HR<1 (favorably prognostic) and 52 ERVs with HR>1 (unfavorably prognostic).

### Prediction of dsRNA formation

To predict potential dsRNA formation from selected ERVs, we utilized ScanFold (v.2.0) [22,23]. ScanFold is a two-stage computational discovery pipeline for RNA secondary structure. The first stage, ScanFold-Scan, utilizes a sliding window algorithm to predict: (1) the minimum free energy (MFE) ΔG°, which predicts the secondary structure and energy of the most stable base pairing arrangement; (2) the ΔG° z-score, which determines if the native MFE ΔG° is more stable than expected by comparing it to those calculated for shuffled versions of the sequence; (3) a p-value for the ΔG° z-score, calculated as the fraction of random ΔG° values more stable than native; (4) the ensemble diversity (ED), which suggests whether the sequence has a propensity of adopting multiple distinct secondary structures (high ED values) or has a propensity of adopting secondary structures which are structurally similar (low ED values). The second stage, ScanFold-Fold, compiles these metrics and generates a list of all stable base pairs and the average metrics observed for each nucleotide of the input sequence. From this list, the most likely arrangement is chosen for each nucleotide until a single consensus model is built for the input sequence. The algorithm selects the arrangement that appears most often for the nucleotide and consistently yields more negative (i.e., more stable) z-scores than other arrangements. For each base pair, the average ΔG° z-score for the most favorable arrangement is calculated and reported (Z_avg_).

Fasta sequences for select ERVs were used as input for ScanFold’s discovery pipeline. The following parameters were used: 120 nucleotide scanning window, 37 ⁰C temperature, global refold, RNAfold algorithm, disallow competition, extract structures from the -2 z-score dot-bracket notation files. ScanFold’s sliding window approach limits predicted interactions to the size of the window; to analyze global ERV structure, each ERV was folded in its entirety using RNAfold alongside 100 sequences produced by shuffling that ERV to calculate the z-score for the entire ERV. The ScanFold pipeline can be run using the scripts available on GitHub (https://github.com/moss-lab/ScanFold2.0) or for use through a web server (https://mosslabtools.bb.iastate.edu/scanfold2).

### In silico immunogenicity and pathways analysis

To assess possible associations between prognostic ERV expression and host immune response, we used the R package EaSIeR to quantify immune cell infiltration based on the quanTISeq algorithm and the following previously published anti-tumor immune hallmark signatures: cytolytic activity (CYT) [3], Roh immune score (Roh_IS) [24], chemokine signature (chemokines) [25], Davoli immune signature (Davoli_IS) [26], IFNy signature (IFNy) [27], expanded immune signature (Ayers_expIS) [27], T-cell inflamed signature (Tcell_inflamed) [27], immune resistance program (RIR) [28] and tertiary lymphoid structures signature (TLS) [29].

For individual gene expression analysis, we used a single-sample version of gene-set enrichment analysis (ssGSEA), which defines an enrichment score as the degree of absolute enrichment of a gene set in each sample within a given data set. ssGSEA analysis was performed on log2(TPM+1) and scaled by pathway z-score, which is the difference between the error-weighted mean of the expression values of the genes in each pathway and the error-weighted mean of all genes in a sample after normalization.

### Statistical analysis

For comparison between two groups, the significance of normal and non-normal distributed variables was estimated using the nonpaired Student’s t-test and Mann-Whitney U test, respectively. For comparisons between more than two groups, one-way analysis of variance (ANOVA) and the Kruskal-Wallis test were used as parametric and nonparametric methods, respectively. Unadjusted p-values were adjusted for multiple hypothesis testing using either Bonferroni or FDR. Statistical significance was set at p<0.05 (two-sided).

### Data visualization and code availability

All raw data processing was conducted using R software (version 4.3.2). All visualizations were performed in R or GraphPad Prism 9 (GraphPad Software, San Diego, CA, United States). Kaplan-Meier plots were drawn using the “ggsurvplot” function in the “survminer” package. The ggplot2 (version 3.4.4) and ggstatsplot (version 0.12.2) packages were used for data plotting [30,31]. Unless otherwise noted, all code and scripts can be found at https://github.com/hchintalapudi/ERV-Project-analysis.

## Results

### Characterization of ERV expression in TCGA cohorts

To advance our understanding of ERV expression in solid tumors, we first assessed global ERV expression in treatment-naive primary tumors using bulk RNA-seq data from TCGA. We selected 8 solid tumor types, which vary in terms of known immunogenicity and response to immunotherapy (S1 Fig). In total, 2,948 samples with clinical data were included. We employed a recently developed pipeline, Telescope, to quantify ERV expression from bulk RNA-seq data [18]. This pipeline estimates the expression of 14,968 ERV elements resolved to specific genomic locations. We defined expression as a minimum of 10 mapped read counts in at least 50% of samples. In total, 7,096 unique ERV transcripts met our expression criteria. On average, 2,736 ERVs were expressed in each tumor type, accounting for 18% of Telescope’s annotated transcripts. Kidney cancers (KIRC) had the highest number of expressed ERVs (n = 3,338), while liver cancers (LIHC) had the lowest (n = 2,229) (Fig 1A).

**Fig 1.**
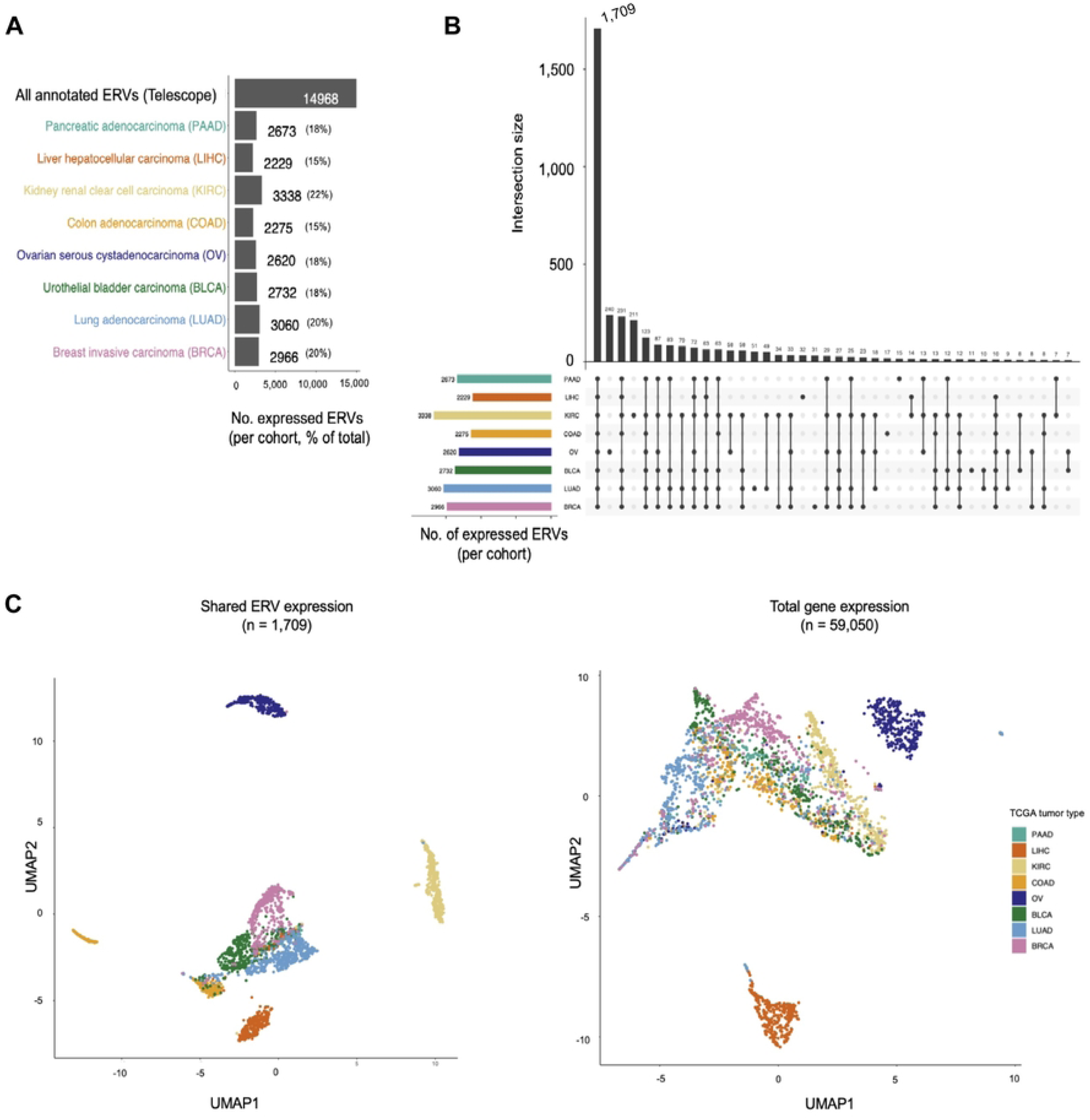
Landscape of ERV expression in 8 types of solid cancers. **A,** Numbers of expressed loci identified in the respective types of cancers. **B,** UpSet plot displaying frequencies of individual ERV loci expressed in one or more TCGA cancers. **C,** UMAP plot showing sample similarity based on normalized expression of shared ERVs (*n*=1,709) or human GRCh38.p12 coding and non-coding genes (*n*=59,050) by multidimensional scaling.

Family-level ERV expression profiles were computed from Telescope’s locus-specific profiles by summing expression across all locations within each subfamily. ERVLE, ERV316A3, MER4, HERVL, HERVH, MER41, and ERVLB4 were the most expressed ERV families across tumors, accounting for over half of expressed ERVs across all cohorts. Additionally, these families were represented in consistent proportions irrespective of tumor type (S2 Fig).

Interestingly, over half of the expressed ERVs in each tumor type were ubiquitously expressed (i.e., shared) across all 8 tumor types (n = 1,709 transcripts) (Fig 1B). The expression profiles of shared ERVs accurately recapitulated both different body sites and tissue types, particularly for kidney and colorectal cancers, which were differentiated more clearly by ERV expression than by coding and non-coding genes (Fig 1C).

### Identification of ERVs associated with survival

To establish whether ERVs are associated with clinical outcome, we generated multivariable univariate Cox proportional hazard models (adjusted for patient age, tumor stage, and tumor grade, when available) to determine whether the expression of each ERV was associated with overall survival (OS) for each tumor type. Thus, each expressed ERV transcript in the corresponding TCGA dataset was tested for a tumor-type-specific survival association. Bladder (BLCA), kidney (KIRC), lung (LUAD), breast (BRCA), and liver (LIHC) cancers all possessed significantly prognostic ERVs after multiple hypothesis correction (FDR<0.25). Of the 7,096 unique ERVs tested, 341 had a favorable association with OS (HR<1), and 2,477 had an unfavorable association (HR>1) (S1 Table). Overall, BLCA and KIRC cancers had the highest number of prognostic ERV transcripts.

Indeed, expression levels of individual ERVs were strongly associated with patient survival in individual tumor types (S3 Fig). For each prognostic ERV (FDR<0.25), samples in each cohort were allocated to groups (high or low) using the ERV’s median expression as a cut-off. In BLCA, KIRC, and LUAD samples, for example, high (i.e., above median) expression of MER4_21q22.3a (HR=0.74), HML5_Xq27.3 (HR=0.89), and Harlequin_1p36.33 (HR=0.76), respectively, identified patients with prolonged survival (S3 Fig).

Next, we sought to identify a set of shared prognostic ERVs in all tumor types (i.e., pan-cancer) using the 1,709 ubiquitously expressed ERVs as input. To reduce the dimensionality of our dataset, we applied a least absolute shrinkage and selection operator (LASSO) penalization to select the representative top predictor ERVs for our pan-cancer survival model (Fig 2A). 449 ERVs were retained after LASSO penalization (see Methods). We applied a multivariable Cox proportional hazards regression model as described above. Of the 449 LASSO-retained ERVs, 94 were significantly prognostic at FDR<0.05; 42 prognostic ERVs were favorable (HR<1), and 52 were unfavorable (HR>1).

**Fig 2.**
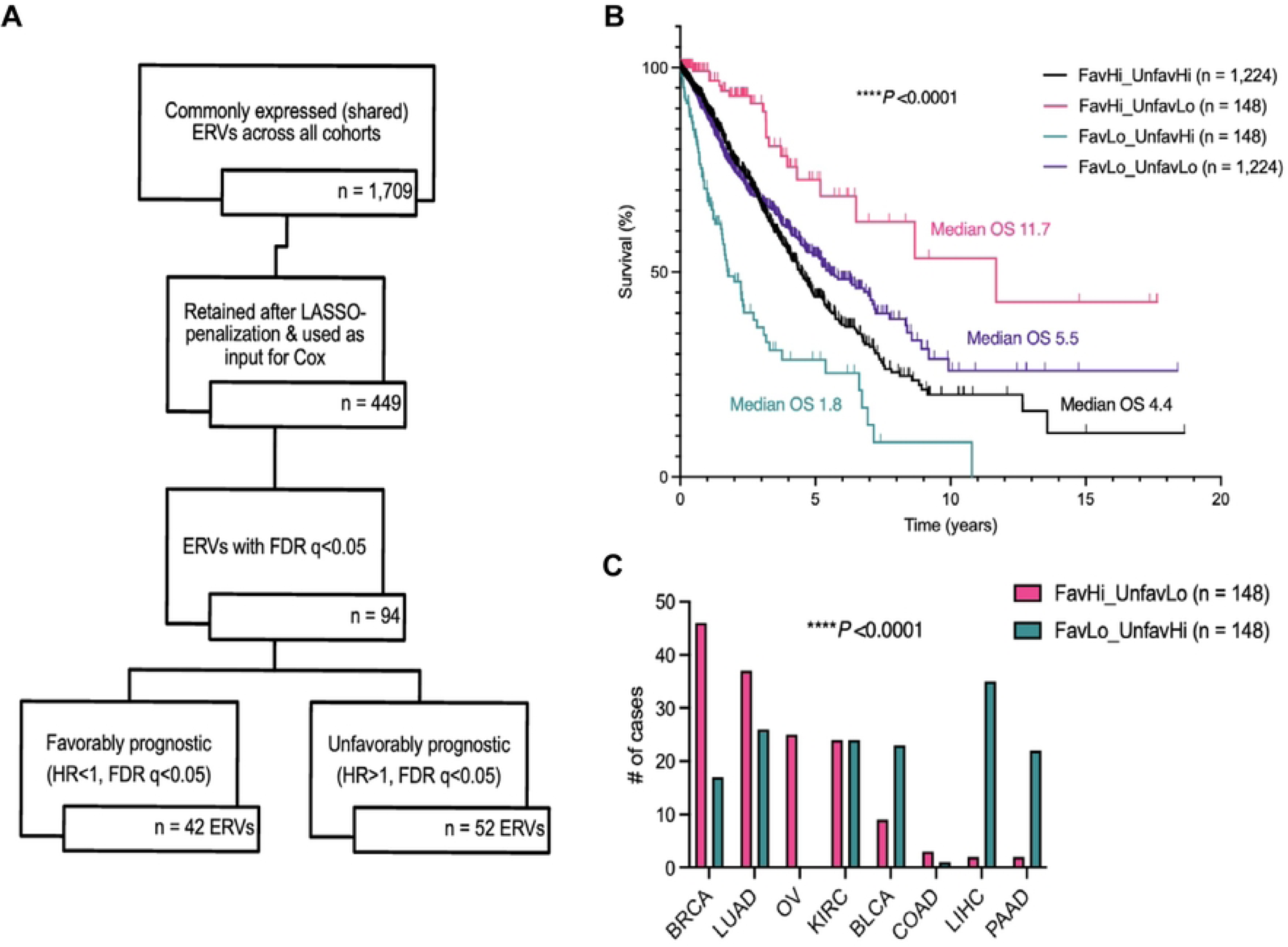
ERV expression is predictive of patient survival. **A,** Schematic representation of the steps and datasets used in deriving a shared prognostic ERV signature across all 8 TCGA tumor types. **B,** Kaplan-Meier curves demonstrating overall survival (OS) of patients stratified according to high (above median) and low (below median) expression of the 42 favorable and 52 unfavorable LASSO-selected ERVs (FDR<0.05). Survival time expressed in years from diagnosis date. P values were obtained from the log-rank Mantel-Cox test. **C,** Chi-squared contingency analysis of tissue-of-origin for the most favorable FavorableERV^Hi^/UnfavorableERV^Lo^ (pink) and least favorable FavorableERV^Lo^/UnfavorableERV^Hi^ (teal) patient groups.

We then classified patients based on favorable and unfavorable ERV expression. We summed up the normalized read counts for each prognostic ERV contained in either the favorable or unfavorable set to generate a favorable and unfavorable prognostic score for each patient, respectively. The median value of the summed normalized read counts within each prognostic set was used as the cut-off into high and low categories. This resulted in 4 patient groups: high expression of both favorable and unfavorable sets of prognostic ERVs (prognostic high), high expression of favorable ERVs with low expression of unfavorable (favorable high), low expression of favorable ERVs with high expression of unfavorable ERVs (unfavorable high), and low expression of both prognostic sets (prognostic low). With these classifiers, we could predict survival outcomes for all 2,948 patients included in this study (Fig 2B). Those with high expression of favorable ERVs but low expression of unfavorable ERVs had the longest survival (median OS 11.7 years). This contrasts patients with high expression of unfavorable ERVs and low expression of the favorable set (median OS 1.8 years). We found that the favorable high group (n=148) was enriched for breast, lung, and ovarian cancer cases, whereas bladder, liver, and pancreas cancer cases were more frequently represented in the unfavorable high group (n=148) (Fig 2C). Nevertheless, associations between patient survival and ERV expression remained evident when broken down by individual tumor type (S1 File). To best capture differences and possible mechanisms by which prognostic ERV expression might influence survival, we focused the remainder of our analyses on the favorable high and unfavorable high patient groups.

### Meta-feature characterization of prognostic ERVs

We explored the genomic meta-features of prognostic ERVs. There were no significant differences in transcript length, coding potential, or chromosomal location between favorable (n=42), unfavorable (n=52), and non-prognostic (n=1,604) shared ERVs (Fig 3A-D).

**Fig 3.**
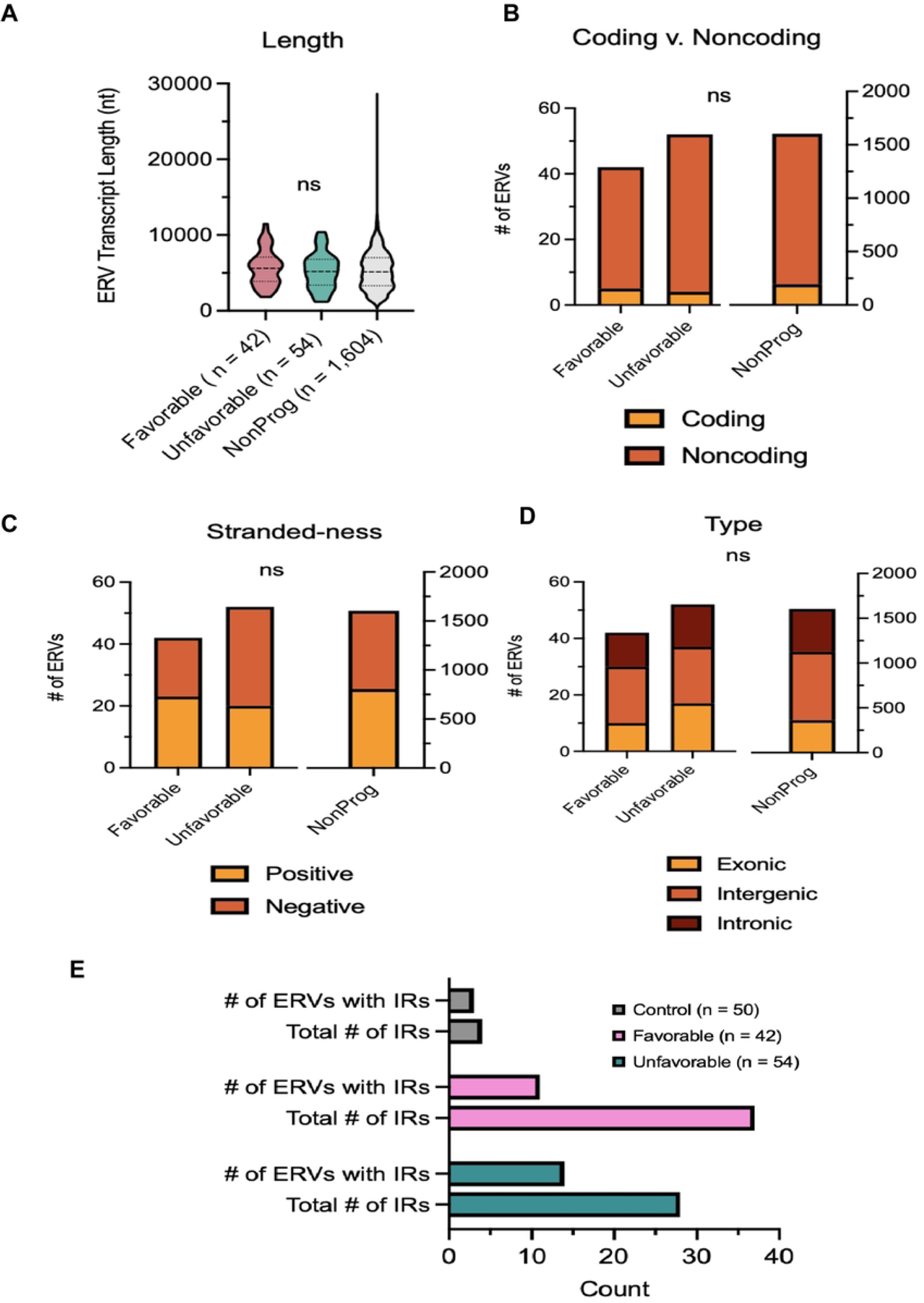
Genomic meta-features between prognostic ERV groups. **A,** Transcript length of favorable (n=42), unfavorable (n=54), and non-prognostic ERVs. **B,** Protein-coding potential of ERVs based on intrinsic sequence composition using CNIT. **C,** DNA strandedness of favorable (n=42), unfavorable (n=54), and non-prognostic ERVs. **D,** Mapped regions with favorable (n=42), unfavorable (n=54), and non-prognostic ERV sequences. **E,** Frequency of inverted repeats (IRs) in favorable (n=42), unfavorable (n=54), and non-prognostic ERV sequences using IR Finder.

We also assessed both prognostic and non-prognostic ERVs according to the presence of inverted repeats (IRs). When a retrovirus integrates into the genome, it often creates inverted repeats – single-stranded nucleotide sequences followed downstream by their reverse complement. This is because the reverse transcription process that converts the viral RNA into DNA can create complementary sequences repeated in reverse order. We found no difference between favorable and unfavorable ERVs concerning the frequency of IR sequences (Fig 3E). We did, however, find a higher quantity of inverted repeat sequences among prognostic compared to non-prognostic ERVs (Fig 3E).

### Prognostic ERVs possess base-pair rich secondary structures with a propensity for ordered stability

In line with our finding that prognostic ERVs are enriched for IR sequences, we hypothesized that these ERV RNAs possess distinctive structural characteristics reflective of their association with patient survival. To predict the secondary structure of these RNA transcripts, we utilized ScanFold (v.2.0) [22,23], a tool that identifies genomic regions with unusual thermodynamic stability and structural conservation. Functional or highly structured RNAs can have a sequence-ordered stability bias, which can be quantified by the thermodynamic z-score [23,32,33]. This score reports the difference between the predicted MFE of folding for a native/ordered RNA and the expected MFE based on the nucleotide content alone. Given the nucleotide sequence composition, a lower z-score indicates lower than expected MFE (more thermodynamic stability). Thus, RNAs with low z-score can have putative evolved/functional structures or extensive ordered folding (e.g., longer-range interactions between IRs) evidenced by unusual stability.

ScanFold predicts RNA secondary structures by first fragmenting input sequences into multiple overlapping windows, where each fragment is folded *in silico* and various thermodynamic properties are calculated. Next, models of structure and predicted ordered biases in RNA structure are combined to generate consensus base pairs weighted by ordered stability. Our initial ScanFold run provided local scans of the folding landscape across our prognostic and non-prognostic control RNAs, quantifying discrete local motifs with a high propensity for ordered structure. We found that favorably prognostic ERVs possessed a more stable secondary structure, as evidenced by significantly lower local z-scores (Fig 4A). This indicates the potential for favorable ERVs to be inherently structured. Indeed, when considering global structure (by using predicted base pairing beyond the 120 nt scanning window to calculate global z-scores), we found significantly negative values whether individual ERVs were across all ERVs, where prognostic ERVs possessed more stable, global structure as compared to control (Fig 4B). This raises interesting questions about the potential roles of globally ordered RNA structures in prognostic ERVs. Interestingly, we also found that prognostic ERVs with lower z-scores had long dsRNA stretches (Fig 4C and S4 Fig). Taken together, we conclude that prognostic ERVs have a greater potential to form dsRNA.

**Fig 4.**
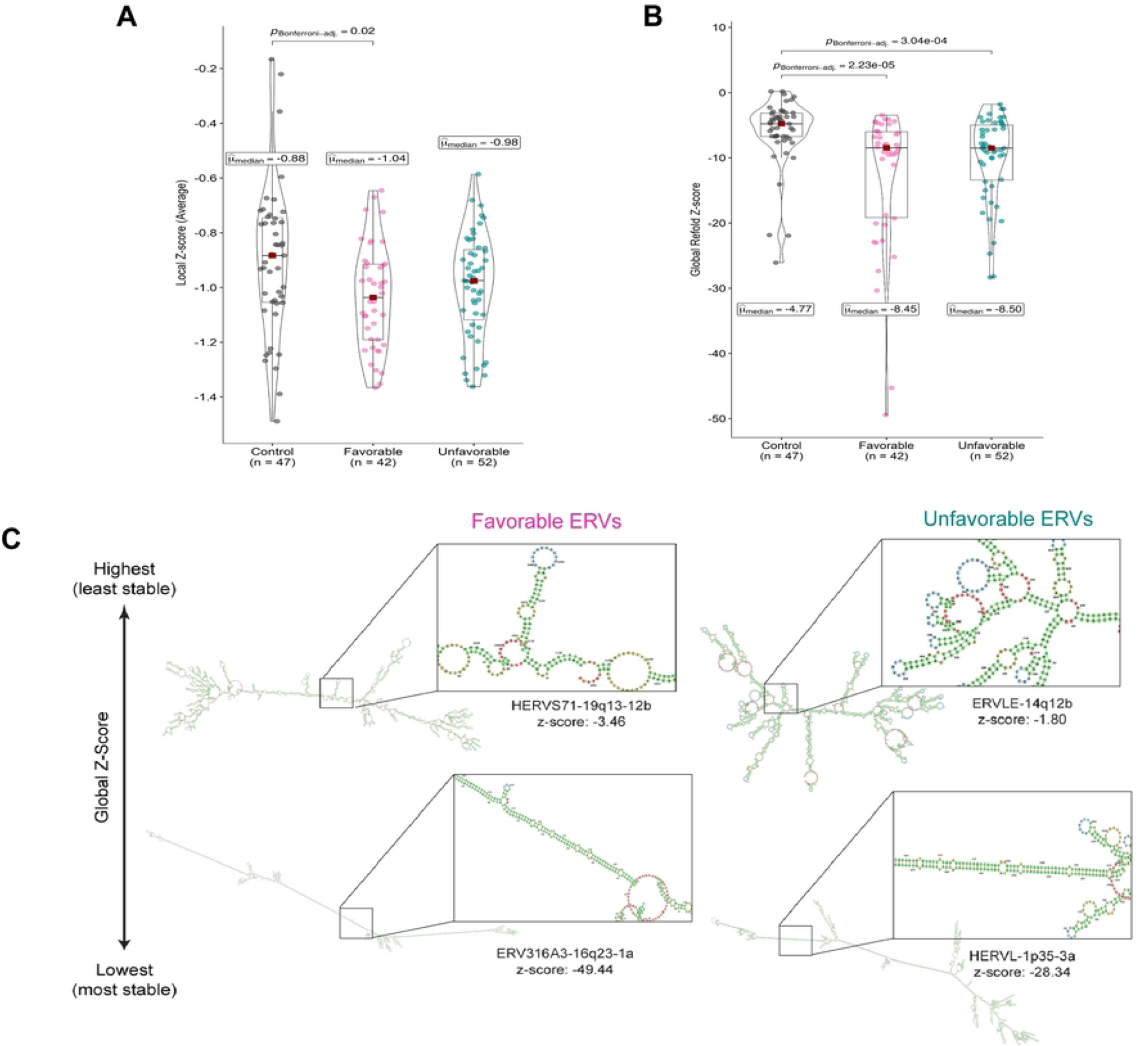
Prognostic ERVs possess unique secondary structures with a propensity for ordered stability. **A,** local RNA structural propensity across groups. Z-score violin plots of each group with a visual trend toward the negative (structured) across prognostic sets. **B,** Global structure. **C,** Predicted secondary structures of select prognostic ERVs. Box and whisker plots: center lines indicate the median; red squares indicate the mean; the box limits indicate the 75th and 25th percentiles; whiskers extend to 1.5 times the interquartile range. Statistical differences between groups were calculated using the Student’s t-test. Data in **A** and **B** are represented as box-violin plots. Center lines (red squares) display the median. *P* values were calculated by Kruskal-Wallis one-way ANOVA with Bonferroni correction for multiple comparisons. Only significant pairwise comparisons are displayed. *\*p < 0.05; **, p < 0.01; ***p < 0.001*.

### Association of prognostic ERVs with expression of immune-related gene sets

Increasing evidence suggests a role for ERVs in the anti-tumor immune response [34–38]. Having identified a set of ERVs whose expression predicts survival, we next investigated the relationship between prognostic ERV expression and transcriptomic signatures of the immune response. We examined the association between prognostic ERV expression and immune signatures using EaSIeR, a tool that derives various quantitative descriptors of the tumor microenvironment, including immune cell composition and tumor immune gene signatures [28].

The first EaSIeR-derived descriptor comprises immune cell fractions obtained with quanTIseq [39]. quanTIseq cell fractions are estimated using a deconvolution approach leveraging cell-type-specific expression signatures for B cells, classically (M1) and alternatively (M2) activated macrophages, monocytes, neutrophils, natural killer (NK) cells, non-regulatory CD4+ T cells, CD8+ T cells, regulatory T (Treg) cells, and myeloid dendritic cells. quanTIseq also provides the fraction of “other” unclassified cells in the mixture, resulting in 11 cellular features. We found a significantly higher proportion of B and CD4+ T cells in the favorable group (Fig 5A). We also saw a higher proportion of Treg cells and M2 macrophages, both immune-suppressive cell types. In contrast, the unfavorable group had a modest but significantly higher proportion of NK cells and neutrophils (Fig 5A).

**Fig 5.**
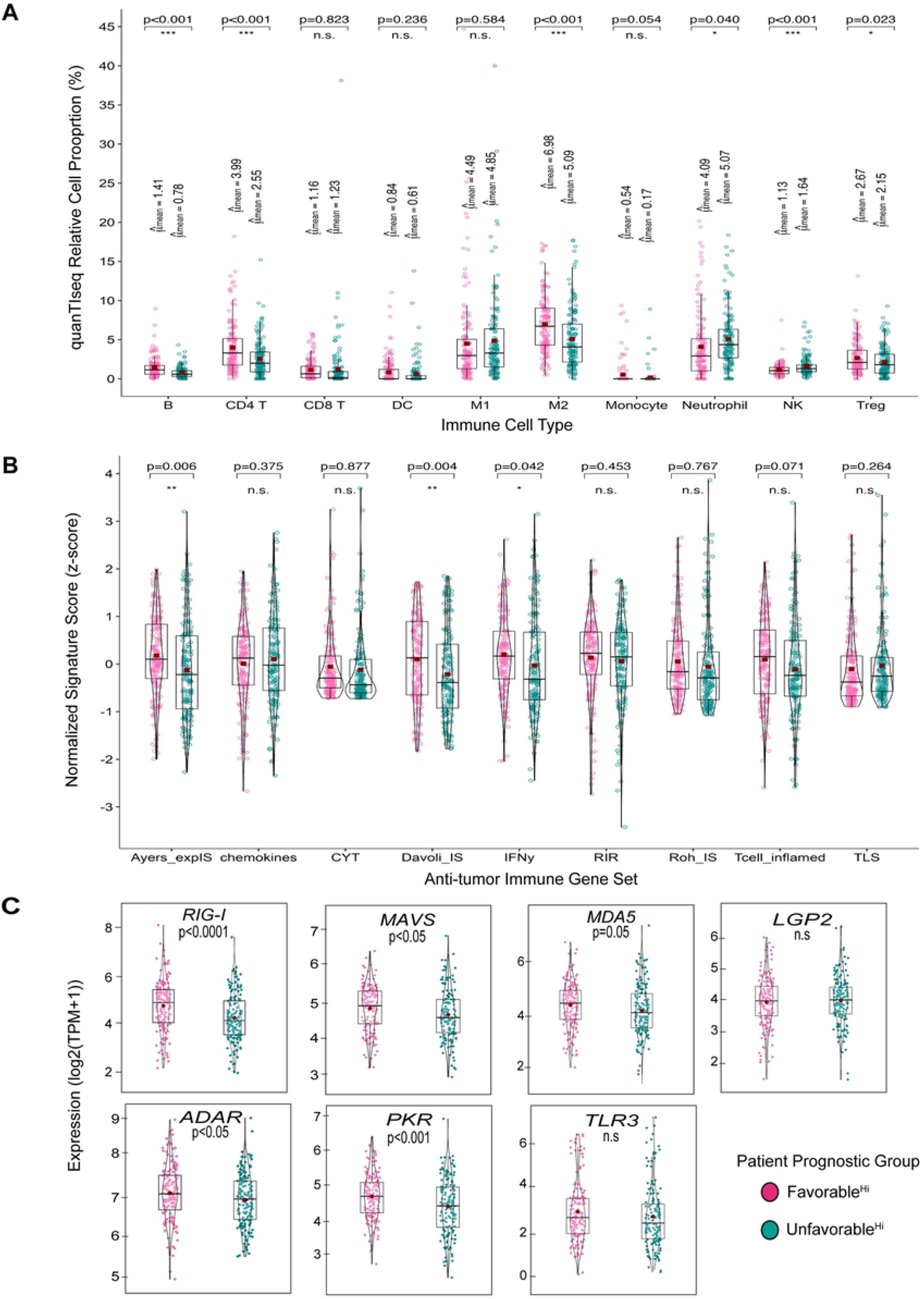
EaSIer transcriptomics-based analysis of the immune response between prognostic ERV groups. **A,** Immune cell type proportion obtained with quanTIseq. quanTIseq absolute scores (i.e., cell type proportions) are relative to the total amount of sequenced cells. **B,** Gene signature scores of previously published anti-tumor immune gene sets. **C,** Expression patterns of innate dsRNA sensor genes. Data is displayed as box-violin plots: center lines indicate the median; red squares indicate the mean; the box limits indicate the 75th and 25th percentiles; whiskers extend to 1.5 times the interquartile range. Statistical differences between favorable and unfavorable groups were calculated using the SStudent’s t-test. Both significant and non-significant (ns) results are shown. *\*p < 0.05; **, p < 0.01; ***p < 0.001.* B, B cells; CD4, non-regulatory CD4+ T cells; CD8, CD8+ T cells; DC, dendritic cells; M1, classically activated macrophages; M2, alternatively activated macrophages; Mono, monocytes; Neu, neutrophils; NK, natural killer cells; Treg, regulatory T cells.

We next used EaSIeR to compute scores for previously published immune response signatures. Three anti-tumor immune hallmark gene signatures were significantly enriched in the favorable group: the Davoli et al. immune signature, the Ayers et al. IFN-γ signature, and the expanded Ayers et al. immune signature (Fig 5B). The Davoli immune signature (Davoli_IS) is derived from the expression of cytotoxic CD8+ T and NK cells. The 6-gene IFN-γ signature contains genes related to IFN-γ signaling (*IFNG, STAT1, CXCL9, CXCL10, IDO1, HLA-DRA*), which have been shown to predict patient response to immune checkpoint therapy in melanoma [27]. The expanded immune signature (Ayers_expIS) extends this analysis by compiling all immune-related genes highly correlated with the IFN-γ signature, including genes associated with cytolytic activity, pro-inflammatory cytokines/chemokines, T cell markers, NK cell activity, antigen presentation, and T cell checkpoints. Despite enrichment for transcripts in the Ayers_expIS gene set in tumors with a favorable ERV expression profile, we did not find a significant difference in the proportion of CD8+ T cells or cytolytic activity between our favorable and unfavorable groups (Fig 5B).

With the propensity of prognostic ERVs to form more stable dsRNA structures, it is possible that ERV-derived dsRNAs activate innate immune sensors, such as toll-like receptor 3 (TLR3) and ubiquitously expressed retinoic acid-inducible gene (RIG)-I-like receptors, triggering antiviral signaling cascades and activating innate immunity. Indeed, we found significantly higher overall expression of individual innate nucleic acid sensors in tumors with high expression of favorable ERVs (Fig 5C). Members of the (RIG)-I-like pattern recognition receptor signaling pathway – RIG-I, melanoma differentiation-associated protein 5 (MDA5), and mitochondrial antiviral signaling protein (MAVS) - were all significantly upregulated in our favorable high-patient group. ADAR and PKR were also significantly upregulated in the favorable group. Other pattern recognition receptors, such as TLR3 and LGP2, showed no significant difference between groups. Taken together, our analysis suggests a possible link between shared prognostic ERVs with a propensity to form more stable dsRNA structures within the tumor microenvironment. These ERVs may influence patient survival through downstream activation of innate immune signaling.

## Discussion

Despite making up a relatively large portion of the genome, ERVs have yet to be extensively studied due to repetitive sequences and a historically small reference set of transcripts [40]. However, this situation continues to evolve with the advancement of deep sequencing and computational tools to improve the capture and quantification of ERV transcripts [41]. Recently, there have been significant advances in the annotation of ERVs and the ability to map these repetitive sequences in RNA-seq data to their most likely genomic origin [18,42,43]. These improvements allow identifying and quantifying specific ERVs associated with genotypes and phenotypes of interest.

Here, we used a recently developed annotation tool, Telescope, to quantify the locus-specific expression of near full-length ERV transcripts representing transcriptional units [18]. Our analysis of ERV expression across multiple TCGA tumor types revealed distinct sets of ERVs whose expression levels were associated with better or worse survival in patients. This corroborates other studies demonstrating tumor-specific expression of distinct ERVs displaying either anti-tumor or tumor-promoting qualities [3,15,44–46]. Furthermore, we identified a set of ubiquitously expressed (i.e., shared) ERVs significantly predictive of survival across all tumor types. Interestingly, these ERVs possessed higher levels of inverted repeats than shared ERVs with no prognostic impact.

Among prognostic ERVs, this higher content of inverted repeats may be informative regarding possible mechanisms by which ERVs may influence outcome. Because of their repetitive sequence content and potential for bi-directional transcription, ERV transcripts are prone to form intra- and inter-molecular nucleic acid structures such as dsRNA. To examine this, we utilized *in-silico* prediction tools, ScanFold and RNAfold, to predict if prognostic ERVs were more likely to form thermodynamically stable secondary structures. Highly structured RNAs, such as the long stems formed from ERV IRs, have sequence-ordered stability bias. Our structural data suggest that prognostic ERVs are more likely to form stable dsRNAs than their non-prognostic counterparts. This was supported by significantly lower global z-scores, indicating more stable global structures due to long-range sequence interactions. However, the differences in z-scores between prognostic and control ERVs could reflect the age of infection or genomic incorporation – where more recently incorporated ERVs possess greater amounts of ordered structure due to maintenance of IR base pairing potential. Over time, the accumulation of mutations in ERVs could disrupt base pairing, reflected in the diminished stability (higher z-scores) observed in non-prognostic control ERVs. Nonetheless, our structural analysis raises interesting questions about the potential roles of globally ordered RNA structure in cancer prognosis.

Given the propensity of our prognostic ERVs to form more stable dsRNA structures, we explored immune-related transcriptional differences between groups. Studies continue to uncover a positive association between ERV transcript levels and innate immune activation, resembling a response against viral dsRNA often observed in virally infected cells [4,5,9,13]. In our study, an association between prognostic ERV expression and evidence of an anti-tumor immune response was intriguing but less than definitive. While tumors with high-level expression of favorable ERVs displayed a higher proportion of B cells and CD4+ T cells, they also displayed higher infiltration by immune-suppressive Treg cells and M2 macrophages. There was no evidence of an increased quantity of CD8+ T cells or effector cytolytic activity, suggesting an unlikely role of our shared prognostic ERVs in eliciting an adaptive immune response. This is further supported by our data showing no difference in coding potential amongst favorable, unfavorable, and non-prognostic ERV transcripts, suggesting that the ability to form neoantigens capable of eliciting an adaptive immune response is not the basis for the observed association between positively prognostic ERVs and patient survival.

Reflecting upon the increased propensity for prognostic ERVs to form secondary structures, we did, however, observe increased expression of several innate dsRNA pattern recognition receptors, including RIG-I/MDA5/MAVS, ADAR, and PKR in tumors with high-expression of favorable ERVs. The cytosolic dsRNA sensor MDA5 preferentially binds to the stem-loop structure formed by IRs to activate viral mimicry. Similarly, protein kinase R (PKR)–mediated cell death is activated by IR stem-loop binding. ADAR signaling is upregulated along with other ISGs following viral mimicry induction and acts as a negative regulator of a dsRNA-mediated IFN response. This negative feedback loop shuts down viral mimicry and PKR activation despite the elevated expression of dsRNAs. Thus, further work is required to parse this highly complex pro- and anti-tumor immune regulation system in ERV-expressing tumors.

In summary, we have identified a common set of structurally distinct ERVs that predict survival in multiple human tumor types. Our analysis of ERV expression across 8 solid tumor types reveals distinct sets of ERVs associated with better or worse survival outcomes. Prognostic ERVs display distinct secondary structures with propensities for ordered stability and are associated with the expression of both positive- and negative-regulators of cellular dsRNA responses. The unique structures encoded by identified prognostic ERVs may modulate cell-intrinsic nucleic acid sensing and innate immunity in a manner that potentially influences cancer patient survival.

## Acknowledgments

This work was supported by the Biostatistics and Bioinformatics Shared Resource of the Dartmouth Cancer Center. The results published here are based on data generated by the Cancer Genome Atlas (TCGA) Research Network (https://www.cancer.gov/tcga); we gratefully acknowledge the contribution of donors and research groups involved in this resource.

## Supporting Information

**S1 Fig.** Schematic workflow of this study.

**S2 Fig.** ERV expression at family-level classification across cohorts.

**S3 Fig.** Kaplan Meier curves for select tissue-type specific ERVs identified from Cox multivariable regression.

**S4 Fig.** Predicated secondary structures for highest (least stable structure), median, and lowest (most stable structure) global z-scores across groups.

**S1 Table.** Tissue-type specific prognostic ERVs expressed in each cohort.

**S1 File.** Kaplan Meier survival curves of shared prognostic ERVs by cohort.

## References

1. Lander ES, Linton LM, Birren B, Nusbaum C, Zody MC, Baldwin J, et al. Initial sequencing and analysis of the human genome. Nature. 2001;409: 860–921. doi:10.1038/35057062

2. Leruste A, Tosello J, Ramos RN, Tauziède-Espariat A, Brohard S, Han Z-Y, et al. Clonally Expanded T Cells Reveal Immunogenicity of Rhabdoid Tumors. Cancer Cell. 2019;36: 597–612.e8. doi:10.1016/j.ccell.2019.10.008

3. Rooney MS, Shukla SA, Wu CJ, Getz G, Hacohen N. Molecular and Genetic Properties of Tumors Associated with Local Immune Cytolytic Activity. Cell. 2015;160: 48–61. doi:10.1016/j.cell.2014.12.033

4. Roulois D, Loo Yau H, Singhania R, Wang Y, Danesh A, Shen SY, et al. DNA-Demethylating Agents Target Colorectal Cancer Cells by Inducing Viral Mimicry by Endogenous Transcripts. Cell. 2015;162: 961–973. doi:10.1016/j.cell.2015.07.056

5. Chiappinelli KB, Strissel PL, Desrichard A, Li H, Henke C, Akman B, et al. Inhibiting DNA Methylation Causes an Interferon Response in Cancer via dsRNA Including Endogenous Retroviruses. Cell. 2015;162: 974–986. doi:10.1016/j.cell.2015.07.011

6. Chuong EB, Elde NC, Feschotte C. Regulatory evolution of innate immunity through co-option of endogenous retroviruses. Science. 2016;351: 1083–1087. doi:10.1126/science.aad5497

7. Topper MJ, Vaz M, Chiappinelli KB, Shields CED, Niknafs N, Yen R-WC, et al. Epigenetic Therapy Ties MYC Depletion to Reversing Immune Evasion and Treating Lung Cancer. Cell. 2017;171: 1284–1300.e21. doi:10.1016/j.cell.2017.10.022

8. Goel S, DeCristo MJ, Watt AC, BrinJones H, Sceneay J, Li BB, et al. CDK4/6 inhibition triggers anti-tumour immunity. Nature. 2017;548: 471–475. doi:10.1038/nature23465

9. Sheng W, LaFleur MW, Nguyen TH, Chen S, Chakravarthy A, Conway J, et al. LSD1 Ablation Stimulates Anti-tumor Immunity and Enables Checkpoint Blockade. Cell. 2018;174: 549–563.e19. doi:10.1016/j.cell.2018.05.052

10. Panda A, Cubas AA de, Stein M, Riedlinger G, Kra J, Mayer T, et al. Endogenous retrovirus expression is associated with response to immune checkpoint blockade in clear cell renal cell carcinoma. JCI insight. 2018;3. doi:10.1172/jci.insight.121522

11. Smith CC, Beckermann KE, Bortone DS, Cubas AAD, Bixby LM, Lee SJ, et al. Endogenous retroviral signatures predict immunotherapy response in clear cell renal cell carcinoma. The Journal of Clinical Investigation. 2018;128: 4804–4820. doi:10.1172/jci121476

12. Solovyov A, Vabret N, Arora KS, Snyder A, Funt SA, Bajorin DF, et al. Global Cancer Transcriptome Quantifies Repeat Element Polarization between Immunotherapy Responsive and T Cell Suppressive Classes. Cell reports. 2018;23: 512–521. doi:10.1016/j.celrep.2018.03.042

13. Cañadas I, Thummalapalli R, Kim JW, Kitajima S, Jenkins RW, Christensen CL, et al. Tumor innate immunity primed by specific interferon-stimulated endogenous retroviruses. Nature medicine. 2018;24: 1143–1150. doi:10.1038/s41591-018-0116-5

14. Segovia C, José-Enériz ES, Munera-Maravilla E, Martínez-Fernández M, Garate L, Miranda E, et al. Inhibition of a G9a/DNMT network triggers immune-mediated bladder cancer regression. Nature medicine. 2019;25: 1073–1081. doi:10.1038/s41591-019-0499-y

15. Kong Y, Rose CM, Cass AA, Williams AG, Darwish M, Lianoglou S, et al. Transposable element expression in tumors is associated with immune infiltration and increased antigenicity. Nat Commun. 2019;10: 5228. doi:10.1038/s41467-019-13035-2

16. Brocks D, Schmidt C, Daskalakis M, Nature … J-H. DNMT and HDAC inhibitors induce cryptic transcription start sites encoded in long terminal repeats. 2017.

17. Lanciano S, Cristofari G. Measuring and interpreting transposable element expression. Nat Rev Genet. 2020; 1–16. doi:10.1038/s41576-020-0251-y

18. Bendall ML, Mulder M de, Iñiguez LP, Lecanda-Sánchez A, Pérez-Losada M, Ostrowski MA, et al. Telescope: Characterization of the retrotranscriptome by accurate estimation of transposable element expression. Plos Comput Biol. 2019;15: e1006453. doi:10.1371/journal.pcbi.1006453

19. Love MI, Huber W, Anders S. Moderated estimation of fold change and dispersion for RNA-seq data with DESeq2. Genome Biol. 2014;15: 550. doi:10.1186/s13059-014-0550-8

20. Guo J-C, Fang S-S, Wu Y, Zhang J-H, Chen Y, Liu J, et al. CNIT: a fast and accurate web tool for identifying protein-coding and long non-coding transcripts based on intrinsic sequence composition. Nucleic Acids Res. 2019;47: W516–W522. doi:10.1093/nar/gkz400

21. Warburton PE, Giordano J, Cheung F, Gelfand Y, Benson G. Inverted Repeat Structure of the Human Genome: The X-Chromosome Contains a Preponderance of Large, Highly Homologous Inverted Repeats That Contain Testes Genes. Genome Res. 2004;14: 1861–1869. doi:10.1101/gr.2542904

22. Andrews RJ, Roche J, Moss WN. ScanFold: an approach for genome-wide discovery of local RNA structural elements—applications to Zika virus and HIV. Peerj. 2018;6: e6136. doi:10.7717/peerj.6136

23. Andrews RJ, Rouse WB, O’Leary CA, Booher NJ, Moss WN. ScanFold 2.0: a rapid approach for identifying potential structured RNA targets in genomes and transcriptomes. Peerj. 2022;10: e14361. doi:10.7717/peerj.14361

24. Roh W, Chen P-L, Reuben A, Spencer CN, Prieto PA, Miller JP, et al. Integrated molecular analysis of tumor biopsies on sequential CTLA-4 and PD-1 blockade reveals markers of response and resistance. Sci Transl Med. 2017;9. doi:10.1126/scitranslmed.aah3560

25. Messina JL, Fenstermacher DA, Eschrich S, Qu X, Berglund AE, Lloyd MC, et al. 12-Chemokine Gene Signature Identifies Lymph Node-like Structures in Melanoma: Potential for Patient Selection for Immunotherapy? Sci Rep. 2012;2: 765. doi:10.1038/srep00765

26. Davoli T, Uno H, Wooten EC, Elledge SJ. Tumor aneuploidy correlates with markers of immune evasion and with reduced response to immunotherapy. Science. 2017;355: eaaf8399. doi:10.1126/science.aaf8399

27. Ayers M, Lunceford J, Nebozhyn M, Murphy E, Loboda A, Kaufman DR, et al. IFN-γ–related mRNA profile predicts clinical response to PD-1 blockade. J Clin Investig. 2017;127: 2930–2940. doi:10.1172/jci91190

28. Jerby-Arnon L, Shah P, Cuoco MS, Rodman C, Su M-J, Melms JC, et al. A Cancer Cell Program Promotes T Cell Exclusion and Resistance to Checkpoint Blockade. Cell. 2018;175: 984–997.e24. doi:10.1016/j.cell.2018.09.006

29. Cabrita R, Lauss M, Sanna A, Donia M, Larsen MS, Mitra S, et al. Tertiary lymphoid structures improve immunotherapy and survival in melanoma. Nature. 2020; 1–5. doi:10.1038/s41586-019-1914-8

30. Wickham H. ggplot2, Elegant Graphics for Data Analysis. R. 2016. doi:10.1007/978-3-319-24277-4

31. Patil I. Visualizations with statistical details: The “ggstatsplot” approach. J Open Source Softw. 2021;6: 3167. doi:10.21105/joss.03167

32. Clote P, Ferré F, Kranakis E, Krizanc D. Structural RNA has lower folding energy than random RNA of the same dinucleotide frequency. RNA. 2005;11: 578–591. doi:10.1261/rna.7220505

33. Babak T, Blencowe BJ, Hughes TR. Considerations in the identification of functional RNA structural elements in genomic alignments. BMC Bioinform. 2007;8: 33. doi:10.1186/1471-2105-8-33

34. Cahn AR, Bhardwaj N, Vabret N. Dark genome, bright ideas: Recent approaches to harness transposable elements in immunotherapies. Cancer Cell. 2022. doi:10.1016/j.ccell.2022.07.003

35. Chen R, Ishak CA, Carvalho DDD. Endogenous Retroelements and the Viral Mimicry Response in Cancer Therapy and Cellular Homeostasis. Cancer Discov. 2021. doi:10.1158/2159-8290.cd-21-0506

36. Jansz N, Faulkner GJ. Endogenous retroviruses in the origins and treatment of cancer. Genome Biol. 2021;22: 147. doi:10.1186/s13059-021-02357-4

37. Stricker E, Peckham-Gregory EC, Scheurer ME. HERVs and Cancer—A Comprehensive Review of the Relationship of Human Endogenous Retroviruses and Human Cancers. Biomedicines. 2023;11: 936. doi:10.3390/biomedicines11030936

38. Müller MD, Holst PJ, Nielsen KN. A Systematic Review of Expression and Immunogenicity of Human Endogenous Retroviral Proteins in Cancer and Discussion of Therapeutic Approaches. Int J Mol Sci. 2022;23: 1330. doi:10.3390/ijms23031330

39. Finotello F, Mayer C, Plattner C, Laschober G, Rieder D, Hackl H, et al. Molecular and pharmacological modulators of the tumor immune contexture revealed by deconvolution of RNA-seq data. Genome Med. 2018;11: 34. doi:10.1186/s13073-019-0638-6

40. Mayer J, Blomberg J, Seal RL. A revised nomenclature for transcribed human endogenous retroviral loci. Mobile DNA. 2011;2: 7. doi:10.1186/1759-8753-2-7

41. Ohtani H, Liu M, Zhou W, Liang G, Jones PA. Switching roles for DNA and histone methylation depend on evolutionary ages of human endogenous retroviruses. Genome research. 2018;28: 1147–1157. doi:10.1101/gr.234229.118

42. Tokuyama M, Kong Y, Song E, Jayewickreme T, Kang I, Iwasaki A. ERVmap analysis reveals genome-wide transcription of human endogenous retroviruses. Proceedings of the National Academy of Sciences of the United States of America. 2018;115: 12565–12572. doi:10.1073/pnas.1814589115

43. O’Neill K, Brocks D, Hammell MG. Mobile genomics: tools and techniques for tackling transposons. Philosophical Transactions Royal Soc B Biological Sci. 2020;375: 20190345. doi:10.1098/rstb.2019.0345

44. Attig J, Young GR, Hosie L, Perkins D, Encheva-Yokoya V, Stoye JP, et al. LTR retroelement expansion of the human cancer transcriptome and immunopeptidome revealed by de novo transcript assembly. Genome Res. 2019;29: 1578–1590. doi:10.1101/gr.248922.119

45. Jang HS, Shah NM, Du AY, Dailey ZZ, Pehrsson EC, Godoy PM, et al. Transposable elements drive widespread expression of oncogenes in human cancers. Nat Genet. 2019;51: 611–617. doi:10.1038/s41588-019-0373-3

46. Zapatka M, Borozan I, Brewer DS, Iskar M, Grundhoff A, Alawi M, et al. The landscape of viral associations in human cancers. Nat Genet. 2020; 1–12. doi:10.1038/s41588-019-0558-9

